# The 6-OHDA Parkinson’s Disease Mouse Model Shows Deficits in Sensory Behavior

**DOI:** 10.1101/2024.06.05.597339

**Authors:** Savannah R. Linen, Nelson H. Chang, Ellen J. Hess, Garrett B. Stanley, Christian Waiblinger

**Affiliations:** Program in Bioinformatics, Georgia Institute of Technology, Atlanta, GA, USA; Wallace H Coulter Department of Biomedical Engineering, Georgia Institute of Technology and Emory University, Atlanta, GA, USA; Departments of Pharmacology and Chemical Biology and Neurology, Emory University, Atlanta, GA USA

## Abstract

Parkinson’s disease (PD) is characterized by the degeneration of dopaminergic (DA) neurons in the substantia nigra pars compacta, leading to dopamine depletion in the striatum and the hallmark motor symptoms of the disease. However, non-motor deficits, particularly sensory symptoms, often precede motor manifestations, offering a potential early diagnostic window. The impact of non-motor deficits on sensation behavior and the underlying mechanisms remain poorly understood. In this study, we examined changes in tactile sensation within a parkinsonian state by employing a mouse model of PD induced by 6-hydroxydopamine (6-OHDA) to deplete striatal DA. Leveraging the conserved mouse whisker system as a model for tactile-sensory stimulation, we conducted psychophysical experiments to assess sensory-driven behavioral performance during a tactile detection task in both the healthy and PD-like state. Our findings reveal a range of deficits across subjects following 6-OHDA lesion, including DA loss, motor asymmetry, weight loss, and varying levels of altered tactile sensation behavior. Behavioral changes ranged from no impairments in minor cases to isolated sensory-behavioral deficits in moderate cases and severe motor dysfunction in advanced stages. These results underscore the complex relationship between DA imbalance and sensory-motor processing, emphasizing the need for precise and multifaceted behavioral measurements to accurately capture the diverse manifestations of PD.

**SIGNIFICANCE STATEMENT:** This study explores sensory-motor aspects of Parkinson’s disease using a 6-OHDA mouse model. Leveraging the mouse whisker system, we reveal diverse deficits in tactile sensation behavior due to dopamine depletion. Our findings emphasize the importance of sensory assessments in understanding the diverse spectrum of PD symptoms.

## Introduction

Parkinson’s disease (PD) is a prevalent neurodegenerative disorder, affecting approximately 1 million individuals in the US annually, with 90,000 new diagnoses reported each year (Willis et al. 2022). Diagnosis is based on identifying common motor symptoms such as tremor, bradykinesia, and muscular rigidity. However, PD patients also experience non-motor impairments, including alterations in olfactory, auditory, tactile, nociceptive, thermal, and proprioceptive perception (Oppo et al. 2020; Jafari, Kolb, and Mohajerani 2020; Kesayan et al. 2015; José Luvizutto et al. 2020; Brim and Struhal 2021). Notably, these non-motor symptoms, often referred to collectively as sensory symptoms, precede the manifestation of motor symptoms by two or more years (Pont-Sunyer et al. 2015), offering a potential window for early diagnostic methods.

In the realm of PD research, animal models have become instrumental in understanding the cellular and circuit mechanisms underlying both motor and non-motor deficits, as well as in the development of therapeutic strategies. The 6-hydroxydopamine (6-OHDA) rodent model, involving the injection of the neurotoxin 6-OHDA into the medial forebrain bundle, is a widely used model that induces dopamine (DA) depletion in the striatum, mimicking Parkinson-like symptoms. Originally designed to study motor deficits (Brooks and Dunnett 2009; Francardo et al. 2011; Lundblad et al. 2004), this model has been extended to investigate non-motor deficits in sensory circuits (Ketzef et al. 2017; de la Torre-Martinez, Ketzef, and Silberberg 2023). However, questions persist regarding how animals with this induced disease model behave and interact with their sensory environment, particularly how stimulus perception and behavioral responses are altered in the DA-depleted state.

Here, we investigate the behavioral aspects of mice trained on a sensory-based detection task in both healthy and Parkinson-like states induced by unilateral 6-OHDA lesioning. Specifically, we leveraged the mouse whisker system for tactile sensory stimulation, a model with well-documented behavioral sensitivity (Petersen 2007) and precise control over stimulus delivery for consistent comparisons across animals. In addition to examining histological changes, motor deficits, and weight changes, we implemented a psychophysical experiment to systematically evaluate performance in a tactile detection task following 6-OHDA lesions, as a complement to conventional measures (Aeed et al. 2021; Branchi et al. 2008). We hypothesized that a detailed analysis of behavioral readouts, combined with physiological markers, would reveal specific alterations in sensory processing and subsequent behavioral responses in the DA-depleted state.

Our findings unveil a spectrum of deficits following 6-OHDA lesion, encompassing DA loss, motor asymmetry, and weight loss, and varying degrees of altered tactile sensation behavior evident in our behavioral detection task. Despite observing substantial DA loss in the majority of mice, those displaying severe behavioral deficits consistently exhibited pronounced impairments across all measures. However, some mice with substantial DA loss exhibited only minor behavioral deficits, while others showed markedly reduced detection performance despite intact motor function, suggesting the presence of an isolated sensory deficit. These results underscore the complexity of DA imbalance and highlight the importance of precise and multifaceted behavioral measurements to effectively model the diverse manifestations of PD.

## Methods

### Experimental model and subject details

All experiments and surgical procedures were performed in accordance with the guidelines approved by the Georgia Institute of Technology Institutional Animal Care and Use Committee and conformed to guidelines established by the NIH. Subjects were 12 male mice (C57/BL/6J, Jackson Laboratories), aged 16-20 weeks at the age of experimentation. Only males were used in this study because the incidence rate of PD is 1.5 times higher in men than in women (Wooten 2004). Animals were housed together in pairs of two per cage (following surgical recovery when applicable) and housed in an inverted light cycle (light: 7pm-7am: dark: 7am-7pm) to ensure experiments occurred during animals’ wakeful periods. Housing temperatures remained at 65-75°F (18-23°C) with 40-60% humidity.

### Head-plate implantation

At least 1 week before experimentation, animals were anesthetized with isoflurane, 5% in an induction chamber, and maintained at 1-3% in a stereotaxic frame. Following anesthetization and analgesia, an incision was made across the length of the skull and the surrounding connective tissues and muscles were removed with a scalpel blade. A ring-shaped (inner radius 5 mm) titanium headplate was then attached to the skull to allow for head fixation during experimentation using MetaBond, a three-stage dental acrylic (Parkell, Inc) (Waiblinger et al. 2022). MetaBond was made on ice, allowed to thicken, applied to the skull, and allowed to set for five minutes to attach the headplate. The skull was then covered with a thin layer of MetaBond followed by a silicone elastomer (Kwik-Cast Sealant, World Precision Instruments) to protect future injection sites. During the procedure animals were placed on a heating pad to keep body temperature stable. Sterile techniques were employed throughout the procedure to minimize chances of infection, and no antibiotics were given. Opioid and non-steroidal anti-inflammatory analgesics were administered (SR-Buprenorphine, SC, post operatively and Ketoprofen, IP, post-operatively). Habituation to head-fixation and training began once animals had fully recovered from head-plate implantation.

### Sensory detection task and training

Behavioral training and testing were conducted using a sensory-driven detection task, utilizing the vibrissa/whisker pathway of the mouse. During behavioral training and testing, water intake was restricted to experimental sessions where animals could earn water to satiety. For two days every week, testing was paused, and animals were given access to water *ad libitum*. Bodyweight was monitored daily and remained relatively stable during constant training. If weight dropped more than ∼5 g, supplementary water was given outside of training sessions to maintain the animal’s body weight. We conducted 1-2 training sessions each day comprising 50-300 trials. All experiments were performed in the dark to ensure no visual identification of the galvo-motor whisker actuator. A constant auditory white noise background noise (70 dB) was produced by an arbitrary waveform generator to mask any sound emission from the galvo-motor whisker actuator. All mice were trained on a Go/No-Go detection task employing a protocol as described previously (Ollerenshaw et al. 2012; Stüttgen, Rüter, and Schwarz 2006; Waiblinger et al. 2018, 2019, 2022). In this task, the whisker is deflected at intervals of 6-8 s (flat probability distribution) with a single pulse (detection target). Trials were sorted into four categories, consisting of a ‘hit’, ‘miss’, ‘correct rejection’, or ‘false alarm’. A trial was considered a ‘hit’ if the animal generated the ‘Go’ response (a lick at a waterspout within 1 s of target onset). The 1 s window of opportunity was chosen as it provides sufficient time for a prompt response, given response latencies of approximately 200–500 ms, while also minimizing the likelihood of random guessing. This restriction ensures that rewards are contingent on sensory-guided behavior rather than chance. If the animal did not lick in response to a whisker deflection, the trial was considered a ‘miss’. Additionally, catch trials in which no deflection of the whisker occurred (A = 0°) were considered a ‘correct rejection’ if the animal did not lick (No-Go). If random licks did occur within 1 s of catch onset however, the trial was classified as a ‘false alarm’. Licking that occurred within a 2 s window prior to stimulus onset was punished by resetting time (time-out) and beginning a new inter-trial interval of 6-8 s, drawn at random from a flat probability distribution. These trial types were excluded from the main data analysis.

### Whisker Stimulation

Whisker deflections were carried out using a calibrated galvo-motor (galvanometer optical scanner model 621OH, Cambridge Technology) as described previously (Chagas et al. 2013). Dental cement was used to narrow the opening of the rotating arm to prevent whisker motion at the insertion site. A single whisker was inserted into the rotating arm of the motor on the right side of the mouse’s face at a 5 mm (±1 mm tolerance) distance from the skin, directly stimulating the whisker shaft and largely overriding bio elastic whisker properties (Waiblinger et al. 2022). All whiskers were trimmed to avoid artifacts generated by the arm touching other whiskers. Across mice, the whisker chosen differed (C1, C2, D1, or D2) but the same was used throughout the experimentation within each mouse. Custom written code in Matlab and Simulink was designed to elicit voltage commands for the actuator (Ver. 2015b; The MathWorks, Natick, Massachusetts, USA). Stimuli were defined as a single event, a sinusoidal pulse (half period of a 100 Hz sine wave, starting at one minimum and ending at the next maximum). The pulse amplitudes used A = [0, 2, 4, 8, 16]°, correspond to maximal velocities: Vmax = [0 628 1256 2512 5023]°/s or mean velocities: Vmean = [0 408 816 1631 3262]°/s and were well within the range reported for frictional slips observed in natural whisker movement (Ritt, Andermann, and Moore 2008; Waiblinger et al. 2022; Wolfe et al. 2008). Figure 2A summarizes the behavior setup, stimulus delivery and logic of the task.

### 6-OHDA lesions

After the collection of behavioral data in the healthy state (pre-lesion, at least 8 sessions of detection testing and one session of rotation), unilateral lesions with 6-OHDA were performed in a subset of animals (6-OHDA, n=10; sham, n=2). The procedures of 6-OHDA injection were adapted from recent studies (Bagga, Dunnett, and Fricker 2015; Ketzef et al. 2017). Animals were anesthetized with isoflurane and mounted on a stereotaxic frame as described in the head-plate implantation section above. Kwik cast was removed, and a small craniotomy was created AP: - 1.2 mm, ML: +1.2 mm, relative to bregma according to the Mouse Brain Atlas in Stereotaxic Coordinates (Frankin and Paxinos 2008). A unilateral injection of 0.2ul of 6-OHDA (15 mg/ml solution 6-OHDA.HBr) was then administered into the left median forebrain bundle at a rate of 0.1ul/min (depth 4.8-5 mm according to the Mouse Brain Atlas in Stereotaxic Coordinates) via a Hamilton syringe (Hamilton Neuros Syringe 700/1700) (Frankin and Paxinos 2008; Thiele, Warre, and Nash 2012). Five-minute waiting periods following both injector insertion and removal were implemented to allow for tissue relaxation. Sham lesioned mice received the correspondent injection protocol with the same volume of vehicle (0.9% saline and 0.02% ascorbic acid). Craniotomies were then sealed with a thin layer of MetaBond and the exposed skull was covered with a silicone elastomer. Opioid and non-steroidal anti-inflammatory analgesics were administered (SR-Buprenorphine, SC, post operatively and Ketoprofen, IP, post-operatively). Animals were allowed to recover for 7–21 days until activity and body weight were normal and then behavioral testing continued.

### Post-lesion care

6-OHDA lesion mice can suffer from a variety of post-operative complications requiring special post-operative care and a continuous health assessment (Masini et al. 2021). After the surgery, animals were placed in single cage and monitored until fully awake. Additional warmth was provided as an infrared light source. In addition to the standard post-operative monitoring and free access to food and water, a maximum of 3ml sterile saline per day was administered SC to encourage rehydration in lesioned mice. Furthermore, lesioned mice received nutritional supplement in form of DietGel Boost (ClearH_2_O) as well as sweetened condensed milk (ClearH_2_O) in food containers on the floor of the cage for 3-10 days (or until body weight stabilized) to increase survival and maintain healthy body weight. The health status of individual mice was assessed daily for a period of 7-21 days post-surgery until activity and body weight were normal. Animals that reached the humane endpoint (defined by the ethical permit) were euthanized.

### Rotation

Unilateral 6-OHDA lesions typically cause highly specific, reproducible rotational measurements and the rotation test is a standard measurement of 6-OHDA induced nigrostriatal pathway lesion and dopamine imbalance in the brain (Brooks and Dunnett 2009). After unilateral median forebrain bundle DA depletion, a postural bias towards the side of the lesion is exhibited. Ipsilateral rotation is driven by an imbalance of DA between hemispheres generating decreased movement on the side of lesion. Mice were placed in a clear plexiglass cylinder (40 cm diameter, 20 cm height) and attached via a flexible harness to an automated counter (Rotation Sensors LE902SR, Container & Sensor Support LE902RP, and Individual Counter LE902CC, Panlab). Ipsilateral and contralateral turns relative to the site of the lesion were recorded for 45 minutes. Turns were counted at 45 degrees and findings were presented as percent left turns, where one turn is equivalent to 360 degrees (8 x 45 degrees). The test was performed for all subjects both prior to and following 6-OHDA (n=10) or sham injection (n=2).

### TH assessment via immunofluorescence

Following experimentation, animals were transcardially perfused using a 4% paraformaldehyde (PFA) solution in phosphate-buffered saline (PBS) (Gage, Kipke, and Shain 2012). Brains were then extracted and post-fixed in PFA for 24 hr before being embedded in OCT for long-term storage at -80°C until sectioning and immunofluorescence. Immunofluorescence staining for tyrosine hydroxylase (TH) was performed as described previously (Ketzef et al. 2017). Sections of 15 uL were collected using a cryostat (CryoStar NX70), rinsed in phosphate buffered saline (PBS) and subsequently incubated for 1 hr in blocking solution consisting of 5% normal goat serum, 0.3% Triton X-100, and 1% BSA in PBS. Following permeabilization, sections were again rinsed in PBS, then incubated with primary antibody, anti-TH rabbit monoclonal antibody (Millipore Sigma, Billerica, MA) diluted 1:500 in BSA-PBS (1%) at 4°C for 24 hr. Sections were then rinsed with PBS and incubated with Cy3-conjugated goat anti-rabbit secondary antibody (Millipore Sigma, Billerica, MA; Jackson Immunoresearch, Philadelphia, PA) diluted 1:800 in BSA-PBS (1%). Following staining, sections were imaged using either a Zeiss 900 confocal microscope (Carl Zeiss, Jena, Germany) at 20x magnification or Leica Stellaris 8 (Leica, Wetzlar, Germany). Images were quantified using ImageJ to assess percent fluorescence change between lesioned and non-lesioned hemispheres.

### TH assessment via high performance liquid chromatography

#### Sample Preparation & Protein Assay

The efficacy of lesioning was assessed using high performance liquid chromatography (HPLC). Mice were sacrificed in the same way as described previously, but no fixation was performed prior to tissue extraction (Gage, Kipke, and Shain 2012). Brain-hemispheres were then separated into non-lesioned (right) and lesioned (left) and placed immediately on dry ice for flash freezing to preserve monoamine concentrations, specifically DA, in the tissue. The tissue was resuspended in 600 uL 0.1 PCA just above freezing temperature and probe sonicated (Branson 450 Digital Sonifier with microtip, Marshall Scientific) on ice with setting 3 and 30% duty cycle. The homogenates were subsequently centrifuged at 4°C at 10,000 x g for 15 min. Supernatants were transferred to 0.22 uM polyvinylidene fluoride polymer (PVDF) microcentrifuge filter tubes and any remaining matter was removed via filtration through spin filter at 800 rpm for 5 minutes. Concentrations of monoamines were acquired through reverse phase HPLC with electrochemical detection. 1000 uL 2% Sodium dodecyl sulfate (SDS) was used to dissolve protein pellets. Quantification of protein was carried out in triplicate 96-well microplates with SpectraMax M5e spectrophotometer (Molecular Devices, Sunnyvale, CA) using the BCA method (Pierce BCA Protein Assay Kit, Thermo Scientific).

#### HPLC Conditions

An ESA 5600A CoulArray detection system equipped with an ESA Model 584 pump and an ESA 542 refrigerated autosampler was utilized to perform HPLC. Separations were performed at 28°C via a Hypersil 150 x 3 mm (3uM particle size) C18 column. The mobile phase consisted of 1.6 mM 1-octanesulfonic acid sodium, 75 mM NaH2PO4, 0.025% triethylamine, and 8% acetonitrile at pH 3. A 25 uL sample was injected and eluted isocratically at 0.4 mL/min. A 6210 electrochemical cell (ESA, Bedford, MA) equipped with 5020 guard cell with potential set at 475 mV was used for sample detection. Analytical cell potentials were –175, 200, 350, and 425 mV. Analytes were identified by matching retention time to known standards (Sigma Chemical Co., St. Louis MO.) and compounds were quantified by comparing peak areas to those of standards on the dominant sensor.

### Experimental timeline

Figure 2B outlines the experimental timeline. Over the course of approximately eight weeks, animals completed an entire set of experiments as described below.

#### Habituation

Habituation to head-fixation began once animals had fully recovered from head-plate implantation for approximately 1 week. Mice were trained to tolerate head-fixation and got accustomed to the experimental chamber for approximately 1 week.

#### Detection Training

Naïve mice received uncued single whisker stimulations in form of a single pulse (A = 16°, P_stim_ = 0.8) interspersed by catch trials or no stimulation (A = 0°, P_catch_ = 0.2). In the early training stage, a water droplet became available immediately after stimulus offset regardless of the animal’s action to condition the animal’s lick response and shape the stimulus reward association. Once animals displayed stable consumption behavior (usually 1-2 sessions), water was only delivered after an indicator lick of the spout within 1000 ms, transitioning the task into an operant conditioning paradigm where the response is only reinforced by reward if it is correctly emitted following stimulus. Learning was measured by calculating the hit p(hit) and false alarm rate p(fa) of successive daily training sessions. A criterion of p(hit) - p(fa) ≤ 0.75 was used to determine successful acquisition of the task. Once an animal reached the criterion, they were considered experts at the task and proceeded into the ‘Detection Testing’ phase.

#### Detection Testing

In this phase, the psychometric curve was measured in repeated daily sessions (1-2 sessions per day) for a minimum of six days. The psychometric curve was measured for all animals using a constant stimuli method entailing the presentation of repeated stimulus blocks containing multiple stimulus amplitudes (A = [0, 2, 4, 8, 16]°). In a single trial, one of multiple stimuli was presented after a variable time interval (6-8 s), each with equal probability (uniform distribution, P = 0.2). A stimulus block consisted of a trial sequence containing all stimuli and a catch trial in a pseudorandom order (e.g. each stimulus is presented once per block). One behavioral session consisted of repeated stimulus blocks until the animal disengaged from the task. Animals had to complete a minimum of n = 50 trials and a session was considered complete once an entire block of stimuli (A = [0, 2, 4, 8, 16]°) was missed. Therefore, trial numbers varied across animals and sessions (n = 50-300 trials). The flexible session length ensures that the data is not affected by the animal’s potential impulsivity or satiation on any given day. Following the six-day testing period, animals were unilaterally injected with 6-OHDA and allowed 7-14 days of recovery. Following recovery animals were again subjected to a ‘Detection Testing’ period which was identical to the one before the lesion to determine the extent of sensory impairments following 6-OHDA lesioning.

#### Length of testing

Over about eight weeks, animals completed a whole set of experiments (Fig. 2b). Note, sessions do not directly correlate with days as two sessions per day were performed in some instances and testing was paused during recovery periods. To resolve the actual time frame of testing, we tracked task performance across days, considering different recovery periods (Fig. 4-1 and table 1). In cases where testing was repeated after longer periods (up to 50 days post-lesion), no differences in behavior was observed. N=2 out of 12 animals reached the humane endpoint likely due to 6-OHDA lesioning and were euthanized before the end of the normal testing period.

**Figure 1.**
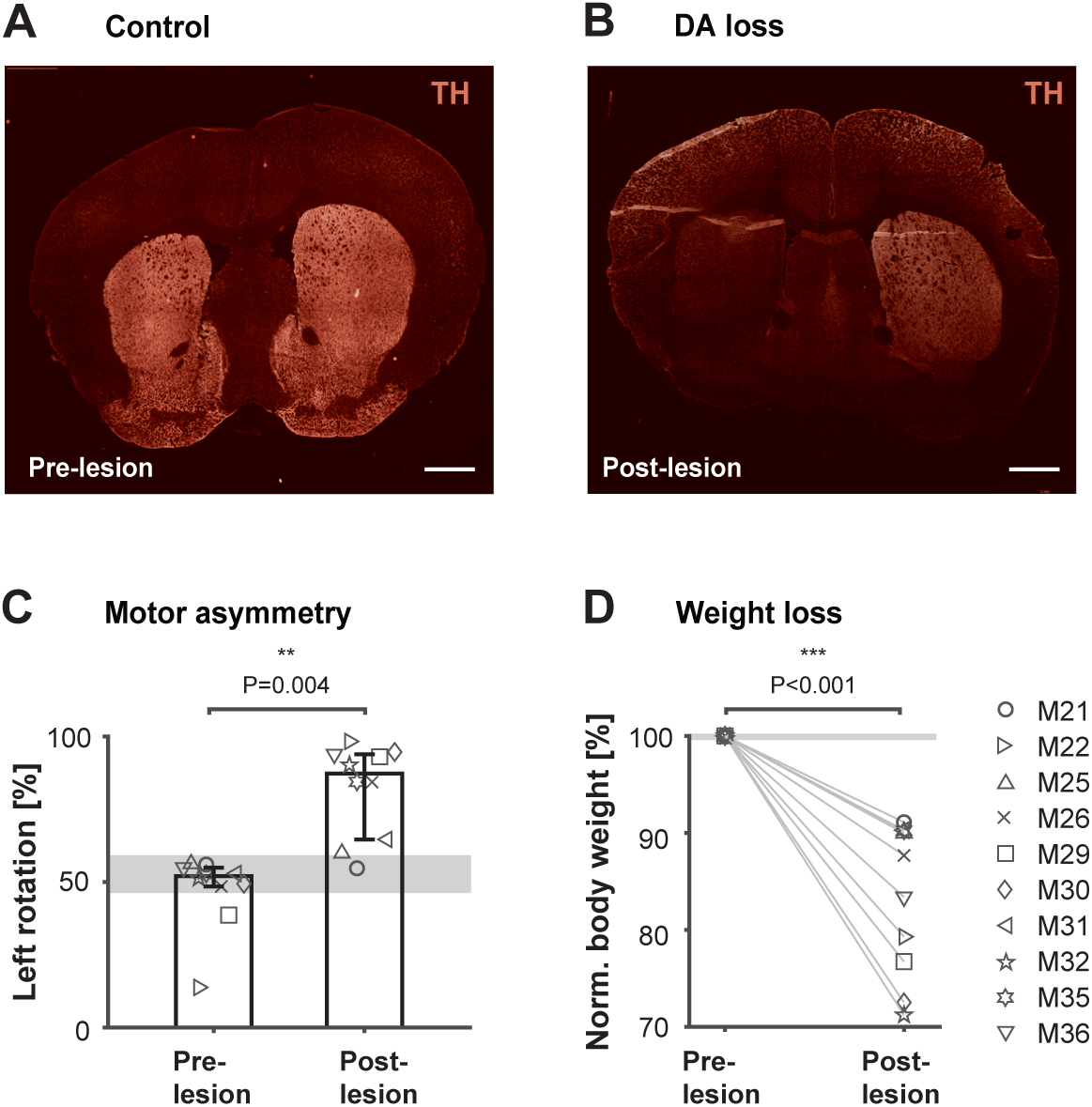
Validation of 6-OHDA mouse model. **A** unilateral injection of 0.2ul of 6-OHDA (15 mg/ml solution 6-OHDA.HBr) was administered into the left median forebrain bundle via Hamilton syringe. Mice were allowed to recover for up to 2 weeks before testing continued. **A**. Coronal sections showing the striatal hemispheres of a control mouse without lesion stained for TH expression. **B**. Coronal section of a 6-OHDA-lesioned mouse brain stained for TH shows over 80% reduction in fluorescence in the dopamine (DA)-depleted hemisphere compared to the control hemisphere. Across all lesioned mice, DA loss was consistently quantified via TH immunofluorescence, showing a median reduction of 77.25% (n = 8, P = 0.008, paired, two-sided Wilcoxon signed rank test). Scale bar: 1 mm. **C**. Percent left turns performed by each mouse in the rotameter test before and after 6-OHDA injection. Bars represent medians across mice with percentiles (n = 10). ** P = 0.004, paired, two-sided Wilcoxon signed rank test. **D**. Body weight before and after 6-OHDA injection. Percent body weight is calculated by identifying the minimum weight within the recovery period. Numbers are normalized to each animals’ weight on the day before the lesion (n = 10). *** P < 0.001, paired, two-sided Wilcoxon signed rank test. Gray areas represent median range across sham operated mice (n = 2).

**Table 1:** Number of sessions and days for each animal pre-lesion, in recovery, and post-lesion.

| Animal ID | Pre-lesion |  | Recovery | Post-lesion |  |
| --- | --- | --- | --- | --- | --- |
|  | Days | Sessions | Days | Days | Sessions |
| 21 | 6 | 6 | 13 | 2* | 3 |
| 22 | 6 | 6 | 13 | 14 | 7 |
| 25 | 6 | 8 | 14 | 15 | 8 |
| 26 | 6 | 10 | 14 | 13 | 10 |
| 29 | 5 | 8 | 23 | 7 | 8 |
| 30 | 7 | 9 | 33 | 3 | 5 |
| 31 | 8 | 8 | 7 | 6 | 13 |
| 32 | 6 | 8 | 15 | 1* | 2 |
| 33 (sham) | 5 | 8 | 19 | 5 | 8 |
| 34 (sham) | 5 | 8 | 19 | 5 | 8 |
| 35 | 6 | 6 | 8 | 7 | 8 |
| 36 | 6 | 6 | 8 | 7 | 8 |

\*Animals euthanized due to humane endpoint.

### Experimental Design and Statistical Analysis

#### Body weight

The weight of each subject was measured daily during detection training, detection testing and recovery phases post surgeries. Percent body weight was calculated by identifying the minimum weight within the recovery period (after 6-OHDA and sham injection). Numbers were normalized to each animals’ weight on the day before the lesion. A paired two-sided Wilcoxon signed rank test was used to assess the significance of the effects of lesions on body weight, comparing measurements from the same animals before and after lesions

#### Rotation test

The rotation test was performed within subjects, prior to and following 6-OHDA (n=10) or sham injection (n=2). Each animal performed the test twice, once before and once after injection. A two-sided Wilcoxon rank-sum test was used to assess the significance of the effects of lesions on rotation, comparing measurements from the same animals in lesioned and healthy conditions.

#### Detection Training

Learning was measured for all subjects by calculating the hit p(hit) and false alarm rate p(fa) of successive daily training sessions. The learning curve was measured by calculating a dprime, d’_behav_, which quantifies the effect size from the observed hit rate and false alarm rate of each training session

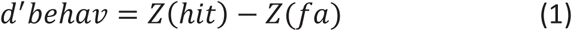

where the function Z(p), p ∊ [0,1], is the inverse of the cumulative distribution function of the Gaussian distribution. A criterion of d’ = 2.3 (calculated with p(hit) = 0.95 and p(fa) = 0.25) was used to determine the end of the detection training period.

#### Detection Testing

The psychometric experiment was performed within subjects, prior to and following 6-OHDA (n = 10) or sham injection (n = 2). The psychometric curve was measured in repeated daily sessions (1-2 sessions per day) for at least six days.

For average psychometric curves across mice (Fig. 2), response probabilities were averaged from all animals that performed the task pre- and post-lesion. For the analysis of individual subjects (Fig. 3), psychometric data was assessed as response-probabilities averaged across sessions within a given stimulus condition.

**Figure 2.**
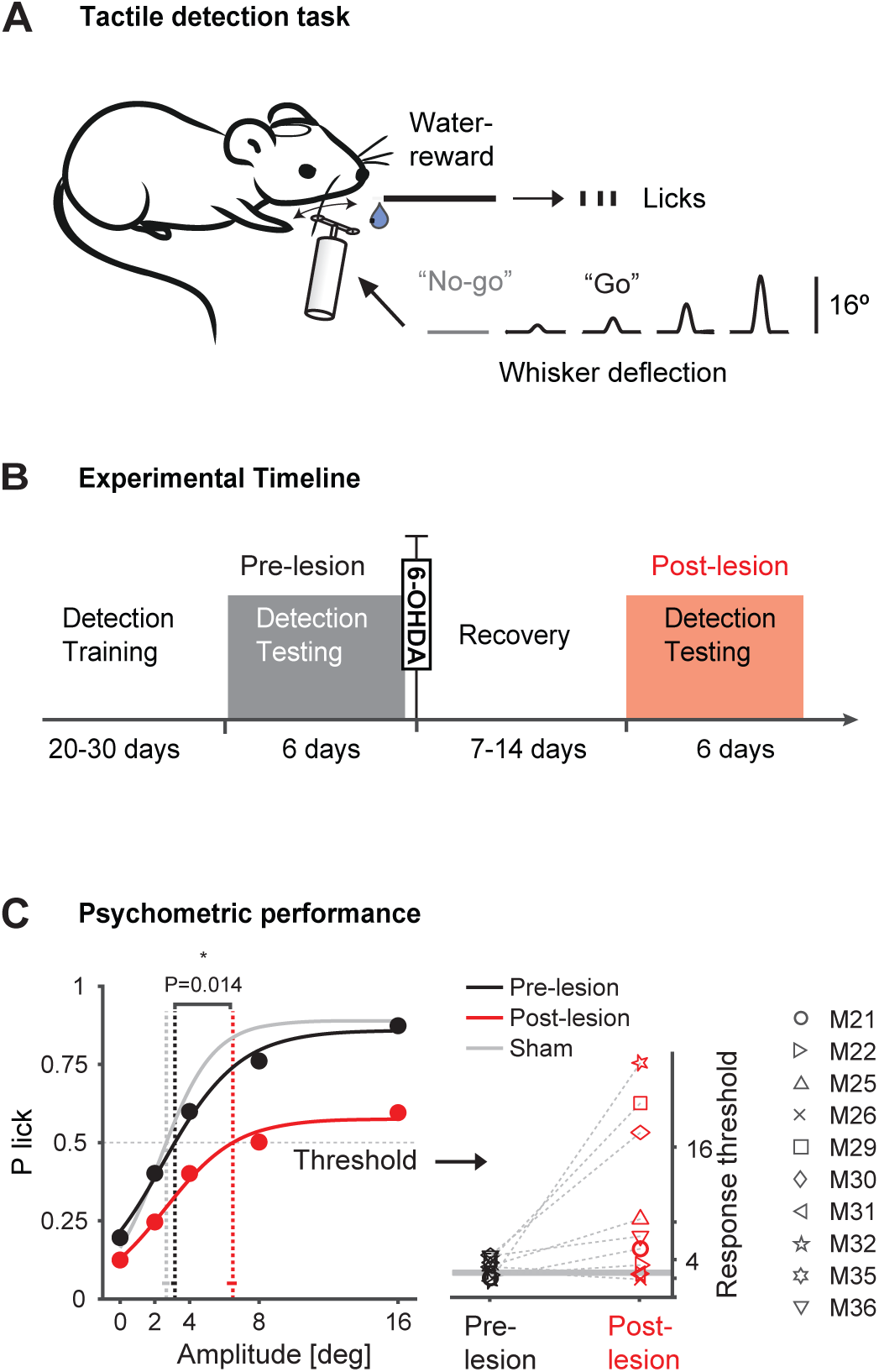
Assessment of sensory capabilities in the 6-OHDA model. **A.** Schematic of the behavior setup, stimulus delivery and behavioral detection task. A punctuate whisker deflection (10 ms) has to be detected by the mouse with an indicator lick to receive reward (”Go”). Licking has to be withheld in catch trials (”No-go”). Reward is only delivered upon correct licks. **B.** Timeline of experiments. Mice were first trained on the detection task. Detection testing was done at least 6 days prior to 6-OHDA injection and repeated after the recovery period. **C.** Left: Psychometric curves and response thresholds before (black) and after 6-OHDA (red) or sham injection (gray). ‘Amplitude [deg]’ represents the degree of whisker deflection. Filled circles correspond to the average response probabilities (P lick) from all animals tested (n=10). Solid curves are logistic fits to the data. Response thresholds are shown as vertical dashed lines with 95% confidence limits at the bottom. * P = 0.014, non-parametric multivariable permutation test for location problem of two independent samples using Fisher’s method for combination of partial tests (preversus post-lesion, n = 10 mice). Right: Response thresholds at P = 0.5 separately shown for all mice before and after the lesion (black and red symbols). Gray bars represent threshold range across sham operated mice.

**Figure 3.**
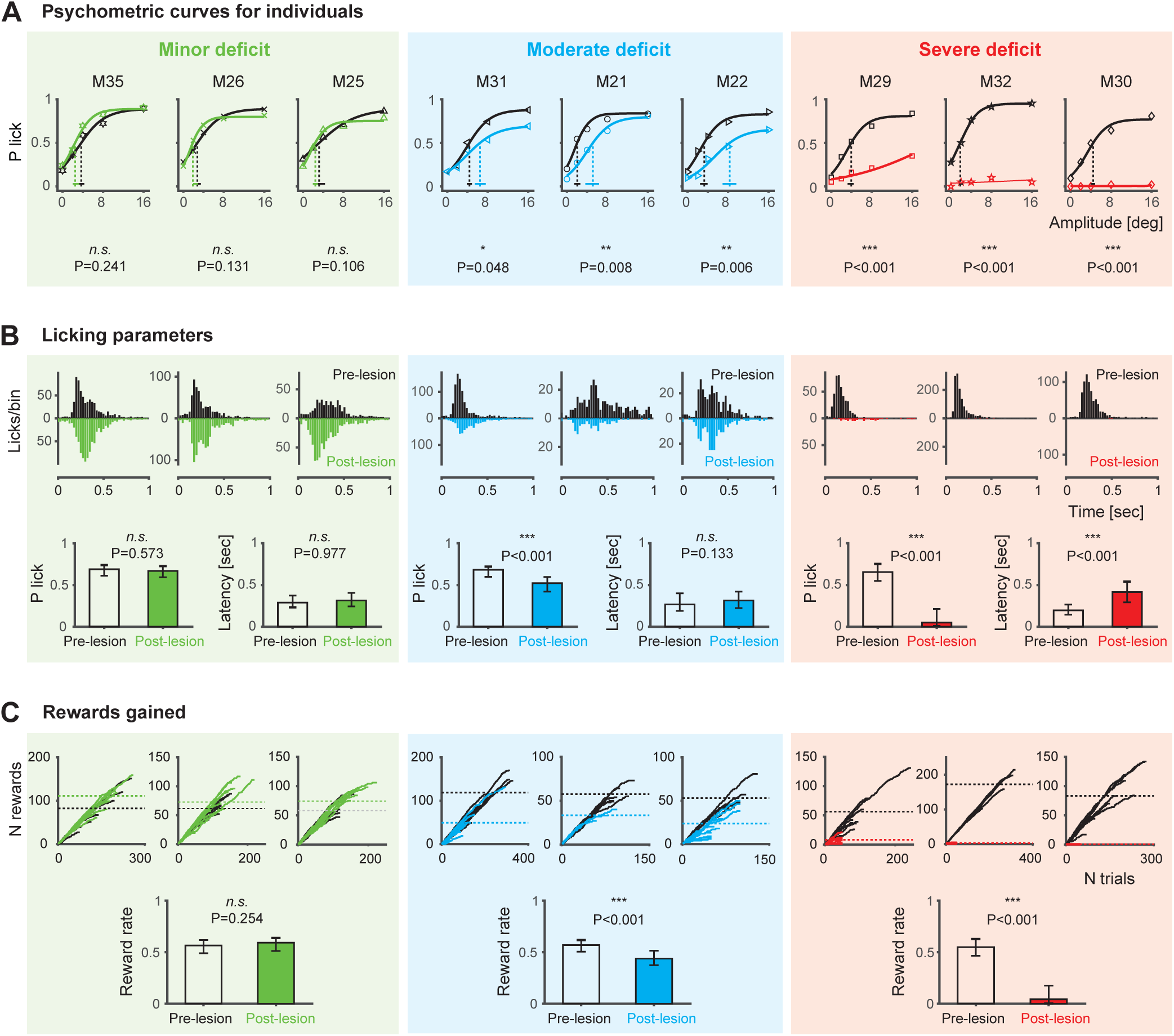
Assessment of sensory capabilities in different individuals. Animals were grouped into 3 categories (Minor, Moderate and Severe deficit) based on severity of psychometric deficits following 6-OHDA lesioning. **A.** Psychometric curves are shown for each animal performing the detection task before (black) and after 6-OHDA lesioning (colored), ordered by performance drop. Points represent response probabilities with different stimulus amplitudes (averaged across sessions), and solid curves are logistic fits. Response thresholds (P = 0.5) with 95% confidence limits are shown as vertical lines. A non-parametric permutation test combined with Fisher’s method was used to compare each subject’s psychometric data before and after the lesion. Mice were then grouped into one of three categories according to the significance level of this test: Minor (*n.s*., P > 0.05), Moderate (P :: 0.05), Severe (P :: 0.001, thresholds >16°, no vertical red lines). **B.** Licking parameters were assessed for animals in each category. Top: Histograms display licks/bin (25 ms bins) for individual mice before and after 6-OHDA lesioning. Bottom: Lick probabilities and latencies were cumulatively calculated for all animals within each category. Bar plots show medians with 25th and 75th percentiles. Statistical differences were assessed using a two-sided Wilcoxon rank-sum test. Lick probability: Minor (*n.s.*, P = 0.573), Moderate (P < 0.001), Severe (P < 0.001). Lick latency: Minor (*n.s.*, P = 0.977), Moderate (*n.s.*, P = 0.133), Severe (P < 0.001). Moderately affected mice show a significant change in lick probability but not in lick latency. **C.** Rewards gained are shown for animals in each category. Top: Number of rewards accumulated across trials by each animal before and after 6-OHDA lesioning. Each line represents a single session, with dashed horizontal lines indicating the average total rewards per session. Bottom: The rate of reward accumulation over trials (slope) was cumulatively calculated for all animals within each group. Bar plots show medians with 25th and 75th percentiles. Statistical differences were assessed using a two-sided Wilcoxon rank-sum test: Minor (*n.s.*, P = 0.254), Moderate (P < 0.001), Severe (P < 0.001).

Psychometric curves were fitted using Psignifit (Frund, Haenel, and Wichmann 2011; Wichmann and Hill 2001). Briefly, a constrained maximum likelihood method was used to fit a modified logistic function with 4 parameters: a (the displacement of the curve), ≤ (related to the inverse of slope of the curve), y (the lower asymptote or guess rate), and A (the higher asymptote or lapse rate) as follows:

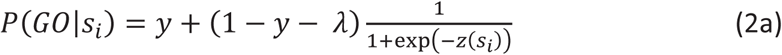

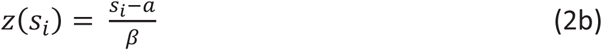

where s_i_ is the stimulus on the i^th^ trial. Response thresholds were calculated from the average psychometric function for a given experimental condition using Psignifit. The term ‘response threshold’ refers to the inverse of the psychometric function at some performance level with respect to the stimulus dimension. Throughout this study, we use a performance level of 0.5.

To assess the effects of the lesion on detection behavior, the psychometric curves and response thresholds were compared in lesion and sham lesion animals, from before and after the injection. Statistical differences between response thresholds were assessed using bootstrapped estimates of 95% confidence limits provided by the Psignifit toolbox. Statistical differences between the full psychophysical datasets were assessed using a non-parametric multivariable permutation test for the location problem of two independent samples, combined with Fisher’s method for partial tests (Arboretti Giancristofaro, Bonnini, and Pesarin 2006; Bonnini, Salmaso, and Solari 2005; Giancristofaro et al. 2010; Pesarin and Salmaso 2010). In the case of Figure 2, the psychophysical data represent the average response probabilities across mice (n = 10) for different stimuli or catch trials, comparing pre- and post-lesion conditions.

#### Impairment severity

The same non-parametric permutation test was applied to each subject’s psychometric data to compare response probabilities for a given stimulus or catch trial across multiple pre-lesion and post-lesion sessions (see Table 1). In the case of Figure 3, subjects were classified into one of three categories based on the significance level of the test results: minor deficit (no significant difference, P > 0.05), moderate deficit (P ≤ 0.05), and severe deficit (P ≤.001). For the severe deficit group, a post-lesion threshold above 16 degrees was used as an additional criterion.

#### Licking and reward parameters

Licking behavior was recorded for each animal before and after the 6-OHDA injection. Lick probability represents the likelihood of the first lick in response to a stimulus within the 1-second ‘window of opportunity,’ excluding subsequent licks related to reward consumption. It serves as a direct metric for determining the psychometric function. In contrast, lick latency refers to the time of the first lick within 1 second after stimulus presentation, reflecting reaction time and motor skills. Reward accumulation was calculated as the cumulative sum of rewards across trials, with each hit trial contributing to a stepwise increase (Figure 3c illustrates trial-by-trial reward accumulation). The slope reflects the rate of reward accumulation over trials, as described previously (Waiblinger et al. 2019). Changes in slope between pre- and post-lesion conditions illustrate the lesion’s impact on reward acquisition. Statistical differences were assessed using a two-sided Wilcoxon rank-sum test

### Data Availability

The data and computer code generated in this study have been deposited in the Zenodo database under accession code https://doi.org/10.5281/zenodo.10733846 and will be made publicly available upon publication of the manuscript.

## Results

This study examines sensory-behavioral deficits in the 6-OHDA mouse model of PD. Using a trained detection task in the mouse vibrissa system, we assessed sensory behavior following unilateral 6-OHDA injections into the medial forebrain bundle (MFB). Relationships among DA loss, motor asymmetry, weight loss, and sensory-behavioral deficits were analyzed to characterize their interactions and contributions to 6-OHDA pathology.

### Validation of the 6-OHDA mouse model

Figure 1 illustrates validation of the 6-OHDA mouse model following unilateral injections into the left MFB. DA innervation in the striatum was assessed using tyrosine hydroxylase (TH) immunofluorescence. In a control mouse, both hemispheres of the striatum appear intact, with no visible difference in fluorescence (Fig. 1A). In contrast, a 6-OHDA-injected mouse exhibits an extensive reduction (81%) in DA innervation in the lesioned hemisphere compared to the non-lesioned hemisphere (Fig. 1B). Across all histologically validated mice, median DA loss was 77.25% (n = 8, P = 0.008, paired, two-sided Wilcoxon signed rank test), with results further supported by high-performance liquid chromatography (92.7%, n = 1).

To further validate DA depletion, all mice underwent a rotameter test before and after 6-OHDA injection (Fig. 1C). This standard test measures 6-OHDA-induced lesions in the nigrostriatal pathway and DA imbalance between hemispheres (Brooks and Dunnett 2009). This DA imbalance leads to a postural bias due to impaired motor abilities on the injection side and is defined here as ‘motor asymmetry’. Prior to 6-OHDA injection, mice exhibited symmetric rotation behavior (median left turns: 52%, n = 10). Following 6-OHDA injection, motor asymmetry emerged, with the majority of mice exhibiting increased rotations toward the lesioned side (median left turns: 87.3%, n = 10, P = 0.004, paired, two-sided Wilcoxon signed rank test). Notably, seven mice displayed clear asymmetry (over 80%), while the remaining three showed a minor deficit comparable to sham controls (approximately 60%). To ensure uniformity of lesioning effects, conventional methods often exclude animals with minimal motor asymmetry (Przedborski and Jackson-Lewis 1995; Ungerstedt and Arbuthnott 1970). In contrast, we conducted a thorough analysis across all animals in this study, including all data to account for potential variability in the effects of 6-OHDA injections among subjects.

In addition to motor asymmetry, we observed significant body weight loss following DA depletion (Fig. 1D), consistent with previous reports in 6-OHDA-treated rodents (Barata-Antunes et al. 2020; Masini et al. 2021). Despite extensive post-operative care, all subjects experienced weight loss within 7–14 days post-injection (median weight loss: 14.47%, n = 10, P < 0.001, paired, two-sided Wilcoxon signed rank test). Notably, weight loss correlated significantly with motor asymmetry, with subjects showing greater weight loss also displaying more pronounced asymmetry (R = 0.81, P = 0.002, n = 10), further validating the efficacy of the 6-OHDA model.

### 6-OHDA lesion affects detection behavior

Twelve mice were trained on a tactile Go/No-Go detection task (Ollerenshaw et al. 2012; Stüttgen, Rüter, and Schwarz 2006; Waiblinger et al. 2018, 2019, 2022), and their performance was evaluated over time in the healthy and PD-like state. Figure 2A depicts the task setup and logic, where mice responded to whisker deflections by either licking a waterspout (Go) or refraining from licking (No-Go) in the absence of a stimulus. Reward delivery depended on correct responses and ‘time-outs’ were used to penalize/discourage impulsive licking. Further details regarding the task are provided in the Methods.

Figure 2B outlines the experimental timeline, with procedures detailed in the Methods section. Briefly, over eight weeks, animals underwent a series of experiments. This included ‘Detection Training’, where animals received single whisker stimulations or no stimulation. Learning progress was assessed daily, and upon meeting criteria, animals proceeded to ‘Detection Testing’. During this phase, the psychometric curve was measured in repeated daily sessions for a minimum of 6 days. Following this testing period, animals underwent unilateral 6-OHDA injection and allowed 7-14 days of recovery. After the recovery period, animals underwent another round of ‘Detection Testing’ post-lesion, mirroring the pre-lesion phase, to assess the extent of behavioral impairments following 6-OHDA lesioning.

Figure 2C shows the averaged psychometric curves for all mice (n = 10) before (black) and after 6-OHDA injection (red), as well as for sham controls (n = 2, gray). Sham-operated mice maintained consistent psychometric performance, indicating that task performance remained stable over time in healthy subjects, ruling out task forgetting or general performance decay. In contrast, lesioned mice exhibited a significant shift in psychometric curve, reflecting a group-level decline in performance (P = 0.014, non-parametric multivariable permutation test; prevs. post-lesion, see methods). In psychophysics, such a shift is typically interpreted as a perceptual deficit (Green and Swets 1966); here, we define it as a ‘sensory-behavioral deficit’, highlighting impaired tactile behavior rather than complete functional loss.

Notably, individual variability in response thresholds was substantial (Figure 2C, right panel), suggesting differential susceptibility to lesion effects or other underlying factors. While this decline is defined as a sensory-behavioral deficit, it is important to consider potential contributions from other task-related variables, such as motor impairments, motivational differences, or overall engagement. These factors were dissected further in the analysis of individual mice (Figure 3) to differentiate motor from sensory components and provide a more detailed understanding of the underlying mechanisms.

### 6-OHDA lesion causes various behavioral deficits across individuals

To explore the behavioral response variability in the 6-OHDA lesioned mice in more detail, we examined multiple behavioral metrics for each animal (Figure 3), including response thresholds, motor lick responses, overall task engagement, and motivation. Statistical comparisons for individual animals were performed using the same non-parametric permutation test applied to the group data. Sham-operated controls exhibited no significant changes in performance (P > 0.05), as shown in Extended Data Figure 3-1, confirming that task performance remained stable over time in the absence of DA depletion. Thus, any observed behavioral changes in the 6-OHDA lesioned mice were likely induced by DA depletion.

Figure 3A illustrates psychometric curves and response thresholds for nine mice performing the detection task in both the healthy (black) and DA-depleted (colored) states (data from one mouse was excluded due to space constraints, though it showed similar trends). Individual analysis revealed varying patterns of task performance following 6-OHDA lesion, enabling us to rank subjects by impairment severity. Based on the shift in psychometric function following the 6-OHDA lesion, animals were categorized into three severity groups: ‘minor’, ‘moderate’, and ‘severe’ deficits (Fig. 3A, from left to right). Mice classified with minor deficits showed no significant performance change (green compared to black curves; P > 0.05). Mice classified with moderate deficits demonstrated a clear performance decline (blue compared to black curves; P ≤ 0.05) but continued to perform the task. In contrast, mice with severe deficits exhibited a substantial performance drop (red compared to black curves; P ≤ 0.001), with post-lesion thresholds exceeding the strongest stimulus intensity, often resulting in task cessation.

To assess for motor dysfunction contributions in psychometric performance, we compared indicator lick probabilities to lick latencies for the minor, moderate, and severe deficit groups. Lick probability represents the likelihood of the first lick in response to a stimulus within the 1-second ‘window of opportunity’. It is a direct metric in determining psychometric function. In contrast, lick latency refers to the time it takes for the animal to produce the first lick, reflecting reaction time and motor skills (Inagaki et al. 2018).

In the severe deficit group, overall task activity drastically decreased, with subjects exhibiting a significant reduction in lick probability and increased lick latency (Fig. 3B, red vs. black histograms and bar plots; P < 0.001, two-sided Wilcoxon rank-sum test). This reduction reflects substantial motor impairments, which may mask potential sensory deficits. Although the performance decline suggests sensory-behavioral deficits, the limited responses in severe cases make it challenging to disentangle motor and sensory contributions. Importantly, these animals continued to exhibit eating, drinking, and grooming behaviors outside the setup, demonstrating some degree of motor control resilience despite task-specific impairments. In the moderate deficit group, lick probabilities were slightly decreased (Fig. 3B, blue vs. black histograms and bars; P < 0.001, two-sided Wilcoxon rank-sum test), while first lick latencies remained normal. This indicates that these animals retained the ability to produce timed motor responses, suggesting intact motor control. However, the observed shift in lick probabilities and corresponding changes in psychometric performance point to a sensory-behavioral deficit. Animals with minor deficits showed no significant changes in lick probabilities or latencies before and after lesioning (Fig. 3B, green vs. black histograms and bars; P = 0.573 and P = 0.977, two-sided Wilcoxon rank-sum test).

Task performance is closely linked to the accumulation of rewards, which reflects the total number of instances in which an animal successfully obtained a reward (Fig. 3C). This measure serves as an approximation of the animal’s motivation, as rewards are a key driver of task performance. We calculated the rate of reward accumulation before and after lesioning. Notably, animals were required to complete a minimum of 50 trials, and a session was deemed complete once an entire block of stimuli (A = [0, 2, 4, 8, 16]°) was missed. The severe deficit group exhibited drastic decreases in reward accumulation (Fig. 3C, red vs. black lines and bar plots, P < 0.001, two-sided Wilcoxon rank-sum test). These severely impaired animals were held in the setup for the minimum number of trials and exhibited very limited activity. In contrast, animals with moderate deficits demonstrated a moderate yet significant decrease in reward accumulation (Fig. 3C, blue vs. black lines and bar plots, P < 0.001, two-sided Wilcoxon rank-sum test) post-lesioning. Finally, animals in the minor deficit group showed comparable reward counts before and after lesioning (Fig. 3C, green vs. black lines and bar plots, P = 0.254, two-sided Wilcoxon rank-sum test).

Our examination of multiple behavioral metrics for individual subjects revealed a spectrum of impairments, reflecting variability in task performance among lesioned animals. We grouped animals based on their task performance post-lesioning into three categories: minor, moderate, and severe deficits. Mice with minor deficits showed normal sensory and motor responses, as indicated by normal psychometric curves, lick probabilities, lick latencies, and reward accumulation. In contrast, mice with severe deficits exhibited gross impairments across all measures. Interestingly, mice with moderate deficits in psychometric function, lick probability, and reward accumulation maintained normal levels of lick latencies. This suggests that, in some cases, sensory impairments in DA-deficient mice may be decoupled from motor dysfunction.

### 6-OHDA lesion causes acute and variable effects over time

Unlike the gradual progression of PD symptoms observed clinically, 6-OHDA induces a swift and complete lesion in the nigrostriatal pathway, commonly injected into the substantia nigra or medial forebrain bundle (Agid et al. 1973; Przedborski and Jackson-Lewis 1995; Tieu 2011). Studies on the motor symptom progression indicate that 6-OHDA-induced lesions can be established with high stability over time (Antony, Diederich, and Balling 2011; Iancu et al. 2005; Quiroga-Varela et al. 2017). However, the progression of sensory-behavioral deficits in this model remains poorly understood. To assess potential changes over time, we systematically analyzed the data from the detection experiments by evaluating behavioral performance across sessions and days in both healthy and PD-like states (Figure 4). Animals were once again categorized into three severity groups (minor, moderate, and severe deficit) as explained in the previous section.

**Figure 4.**
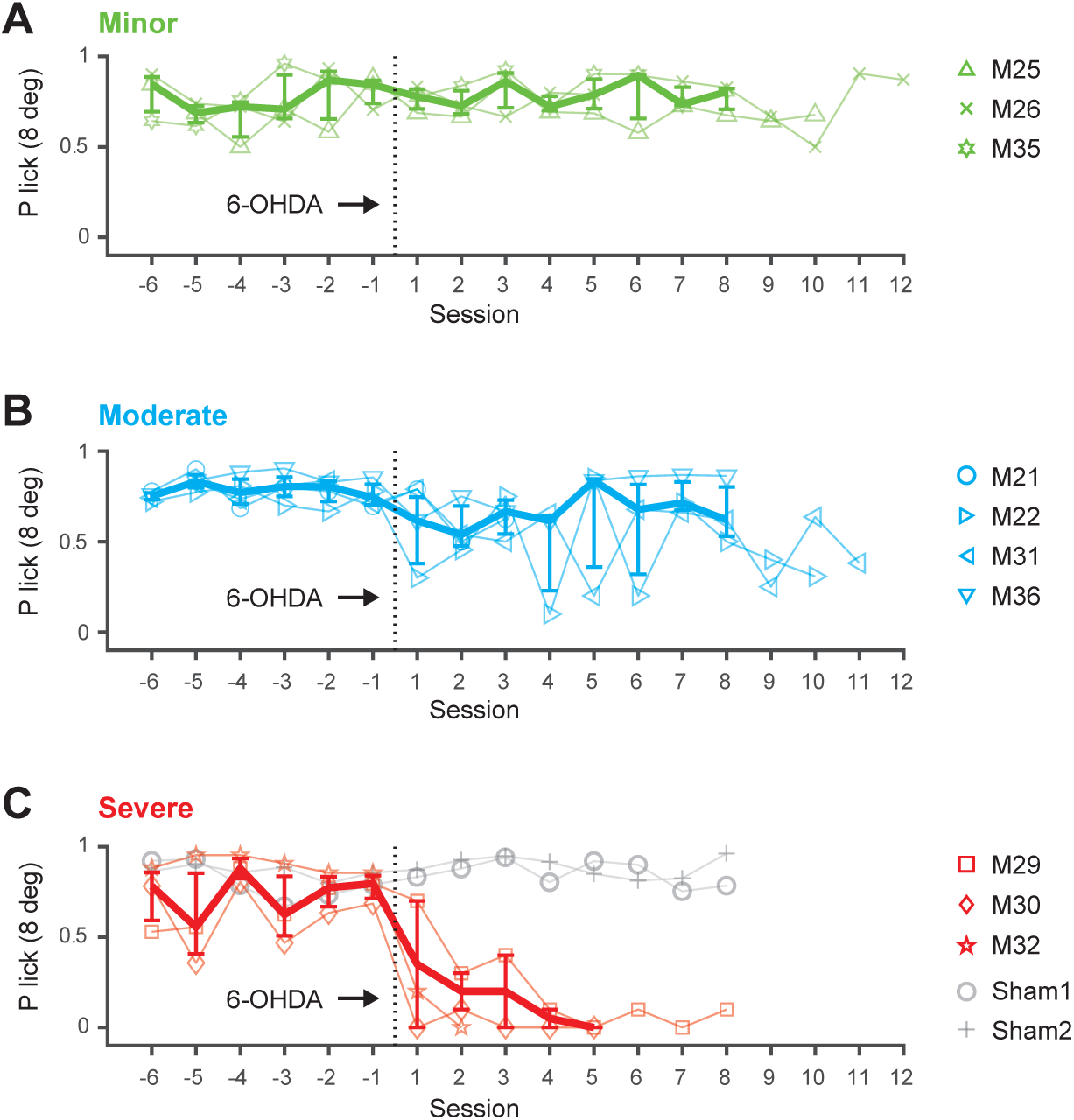
Acute and variable effect in the 6-OHDA mouse model. The 3 categories, minor, moderate and severe, are based on the difference in thresholds from the psychometric curves. **A.** Detection performance (P correct lick with 8 degree whisker deflection) over time (sessions) for animals with minor deficits, before and after the 6-OHDA lesion. **B.** Performance for animals with moderate deficits. **C.** Performance for animals with severe deficits. Symbols represent individual mice. Thick lines represent medians across mice with percentiles (n=3-4). Gray lines and symbols represent data for sham operated mice (n=2).

Figure 4A illustrates the detection performance over multiple sessions for three animals with minor deficits, both before and after the 6-OHDA lesion. Performance is represented as the probability of a correct lick with a salient stimulus (8-degree whisker deflection) across 6 prelesion sessions and up to 12 post-lesion sessions for each animal. Before the lesion, animals exhibited low variability in performance. After the lesion, performance remained unchanged, with consistent levels observed across sessions, suggesting that the sensory capabilities linked to the task likely remained intact in these mice.

Figure 4B illustrates the detection performance over time for four animals exhibiting moderate deficits. While some individuals showed an initial decline in performance shortly after the lesion, noticeable variability emerged across sessions, occasionally within a single day. The session-to-session variability underscores the complexity of the animals’ responses, potentially elucidating the coexistence of sensory and motivational deficits alongside preserved motor capabilities within the detection task, as previously observed for this group of mice (Figure 3, blue panels).

Figure 4C illustrates a substantial decline in detection performance in three severely affected animals shortly after the 6-OHDA injection, evident within the initial two sessions. Following the lesion, performance dropped to nearly zero levels with minimal session-to-session variability. This pronounced and persistent change confirms previous observations that severely impaired animals exhibit very limited activity in response to DA depletion.

Notably, recovery periods varied among the three groups of mice, resulting in differences in the data presented regarding the lesion timeframe. Additionally, due to occasional repetition of sessions within a day and pauses on weekends, sessions do not directly correspond to days. To address this, performance was tracked across days in a separate analysis, considering these variations (Figure. 4-1 and Table 1). The day-based analysis is consistent with the session-based analysis, confirming that task performance did not recover or degrade over time within the measured timeframe.

### Selective correlations between 6-OHDA pathology and behavior

Unilateral 6-OHDA injection induces pathological changes that can be assessed through a combination of direct and functional measures. DA depletion and weight loss serve as direct markers of the lesion’s impact, while motor asymmetry and sensory-behavioral deficits reflect its functional consequences. This section integrates all metrics assessed to explore their relationships and the broader effects of 6-OHDA pathology, as shown in Figure 5.

**Figure 5.**
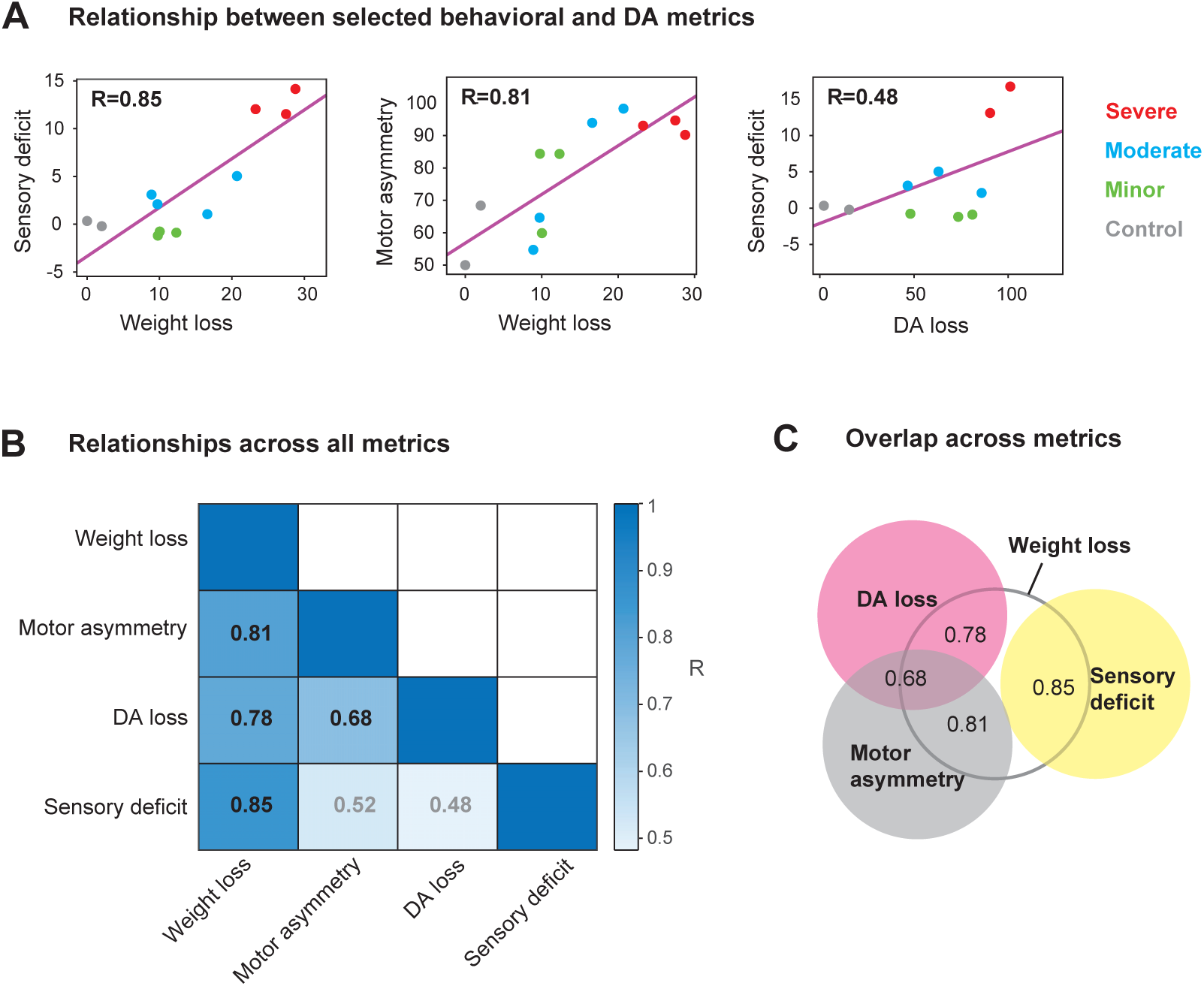
Correlations between pathology and behavior in the 6-OHDA mouse model. **A.** Scatter plots of selected behavioral and pathological metrics. Each point represents data from an individual animal, with colors representing the severity classification. Pearson’s correlation coefficients (R) and least-squares reference lines (magenta, slope = R) are shown. Weight loss strongly correlates with sensory-behavioral deficits (R = 0.85, left) and motor asymmetry (R = 0.81, middle). In contrast, DA loss does not correlate with sensory-behavioral deficits (R = 0.48, right). **B.** Pairwise correlation coefficients (R-values) between all metrics. Significant correlations are highlighted in black ([R = 0.81, P = 0.002]; [R = 0.78, P = 0.008]; [R = 0.68, P = 0.029]; [R = 0.85, P :: 0.001]), and non-significant correlations are depicted in gray ([R = 0.52, P = 0.082]; [R = 0.48, P = 0.158]). P-values are based on t statistics with n-2 degrees of freedom. **C.** Venn diagram summarizing the overlap and distinct contributions of behavioral and pathological metrics.

Figure 5A illustrates representative relationships between key metrics using scatter plots. Weight loss strongly correlated with sensory-behavioral deficits, quantified as changes in response thresholds post-6-OHDA lesion, with greater weight loss associated with more severe deficits (R = 0.85, P ≤ 0.001, left). It also positively correlated with motor asymmetry, with higher weight loss linked to greater asymmetry post-6-OHDA injection (R = 0.81, P = 0.002, middle). In contrast, DA loss showed a weaker, non-significant correlation with sensory-behavioral deficits (R = 0.48, P = 0.158, right) and exhibited substantial variability across subjects, with some mice showing extensive DA loss (> 70%) but only minor to moderate deficits.

Figure 5B and C provide an overview of these relationships. Figure 5B presents the pair-wise correlation coefficients between the various metrics, showing that direct markers like weight loss and DA loss correlate with each other but show inconsistent alignment with functional markers such as motor asymmetry and sensory-behavioral deficits. Weight loss, however, correlated significantly with all other metrics (P ≤ 0.05), illustrated conceptually in the Venn diagram in Figure 5C. Notably, mice with severe sensory-behavioral deficits consistently showed impairments across all measures, while minor and moderate deficits exhibited greater variability, indicating heterogeneity in the effects of the lesion.

Together, these findings establish weight loss as a reliable marker correlating with the physiological and functional impacts of 6-OHDA lesions, spanning sensory-behavioral and motor impairments. The lack of a significant correlation between DA loss and sensory-behavioral deficits underscores individual variability and highlights the complexity of 6-OHDA pathology. Leveraging a sensory-based detection task in this mouse model was pivotal in isolating sensory deficits from motor impairments, reinforcing its utility in uncovering the specific effects of DA depletion on sensory processing and behavior.

## Discussion

While PD is primarily diagnosed based on motor symptoms, sensory deficits are increasingly recognized as an early and underexplored aspect of the disease (Juri, Rodriguez-Oroz, and Obeso 2010; Konczak et al. 2012; Prätorius, Kimmeskamp, and Milani 2003; Richardson and Sussman 2019). Studies have documented impairments across multiple sensory modalities, including olfactory, auditory, tactile, nociceptive, thermal, and proprioceptive processing (Oppo et al. 2020; Jafari, Kolb, and Mohajerani 2020; Kesayan et al. 2015; José Luvizutto et al. 2020; Brim and Struhal 2021). These deficits suggest that PD extends beyond motor dysfunction, affecting sensory processing and behavior, and offering a potential window for earlier diagnosis and intervention. However, the mechanisms underlying sensory and behavioral changes remain unclear, partly due to challenges in isolating sensory components from motor dysfunction in human studies.

To address this gap, we employed the 6-OHDA mouse model, a widely used tool for studying PD-like states. This model induces targeted dopamine (DA) depletion, enabling precise experimental control to dissect the effects of DA loss. Although it lacks the progressive nature of human PD and primarily induces rapid, selective DA degeneration, the 6-OHDA model reliably replicates key aspects of PD-related pathology (Bagga, Dunnett, and Fricker 2015; Francardo et al. 2011; Masini et al. 2021). This experimental setup provides a robust platform for investigating how DA loss affects sensory processing and behavior, complementing human studies where complexity and variability often hinder mechanistic insights.

Using this model, we focused on sensory-behavioral deficits in a psychophysical whisker-based detection task. The mouse whisker system, as a primary sensory modality for rodents, offers well-defined topographic organization, rapid signaling dynamics, and compatibility with controlled behavioral tasks (Petersen 2007), making it ideally suited for linking sensory deficits to DA loss. While we expect similar results in other modalities, such as forepaw stimulation, visual, or auditory systems, these may offer less precise control of experimental conditions.

Our findings revealed a spectrum of behavioral changes, ranging from no impairments in minor cases to isolated sensory-behavioral deficits in moderate cases, and severe motor dysfunction in advanced cases. One consideration in interpreting these findings is the challenge of disentangling sensory impairments from motor dysfunction, as our behavioral task relies on a motor response for readout. While the maintenance of normal response latencies in moderate cases suggests that sensory deficits can be decoupled from motor impairments, this overlap complicates the interpretation of sensory-behavioral changes. In addition, reduced rewards and variable performance in moderate cases point to motivational changes as another factor influencing behavior. Addressing these complexities will require further exploration of the underlying mechanisms to better understand how DA depletion affects sensory and motor interactions.

The variability observed in behavioral outcomes likely reflects a combination of factors, including lesion accuracy, neuronal susceptibility, pharmacokinetics, and individual differences in behavioral responses. While this study focused on overall DA depletion, future research is needed to systematically evaluate lesion placement and its impact on specific striatal subregions. Notably, we found no direct correlation between histologically quantified DA loss and behavioral severity; all mice exhibited substantial DA depletion regardless of their deficits. This suggests that a threshold level of DA loss may need to be surpassed before measurable impairments emerge. Beyond this threshold, compensatory mechanisms or other pathological changes may contribute to the variability in outcomes.

Weight loss emerged as a consistent metric across DA loss, motor asymmetry, and sensory-behavioral deficits. Although the mechanisms differ in timing and progression between our toxin-based model and clinical PD, weight loss reflects acute DA-related pathological effects in the former (Masini et al. 2021) and chronic dopaminergic dysfunction affecting appetite, apathy, and sensory processing in the latter (Aiello, Eleopra, and Rumiati 2015; Cumming et al. 2017; Ma et al. 2018). This parallel highlight its value as an integrative marker for pathology and behavior.

The potential mechanisms underlying sensory deficits in DA-depleted states likely involve disruptions in the basal ganglia-cortex circuitry, with DA playing a critical role in modulating sensory signal processing. Within this loop, cortico-striatal inputs from sensory and motor cortex distinctly influence striatal activity and behavior (Lee et al. 2019), while DA signals in the dorsomedial striatum encode contralateral sensory stimuli and support stimulus-action associations (Moss et al. 2021). DA loss significantly alters striatal sensory responses, disrupting sensory processing (Ketzef et al. 2017; de la Torre-Martinez, Ketzef, and Silberberg 2023). Ketzef et al. demonstrated that DA depletion disrupts laterality coding, in response to bilateral whisker stimulation. Our findings complement this work by incorporating behavioral tasks, revealing sensory-behavioral deficits during unilateral whisker detection. Together, these studies suggest that DA depletion broadly impacts sensory processing, affecting both bilateral and unilateral stimulation. Combining behavioral tasks with electrophysiological recordings and comprehensive stimulation protocols may provide deeper insights into the specific mechanisms of these disruptions.

In summary, our study highlights the utility of the 6-OHDA model for investigating sensory-behavioral deficits relevant to early PD symptoms. The whisker-based detection task proved effective for uncovering hidden variables and assessing different stages of impairment. Our severity-based classification system provides insights into early PD detection and the versatility of the 6-OHDA model. Building on this foundation, research using progressive models, longitudinal measurements, and integrated physiological approaches could further illuminate the complex relationship between sensory processing and DA loss, advancing diagnostic and therapeutic strategies.

## Acknowledgements

S.R.L. was supported by a McCamish Parkinson’s Disease Innovation Program Blue Sky grant. E.J.H was supported by a McCamish Parkinson’s Disease Innovation Program Blue Sky grant and NIH R01NS124764. N.H.C., G.B.S and C.W. were supported by NIH Brain Grants RF1NS128896 and R01NS104928. This study was supported by the Emory University Emory Integrated Cellular Imaging Core Facility (RRID:SCR_023534), with special thanks to April Reedy for her advice on experimental design, image acquisition and statistical help. We thank the Emory HPLC Bioanalytical Core (EHBC), which was supported by the Emory University School of Medicine and the Georgia Clinical & Translational Science Alliance of the National Institutes of Health under Award Number UL1TR002378. Additionally, we thank Aqua Asberry from the Georgia Institute of Technology’s Research Histology Core for her assistance in specimen embedding and cryostat sectioning.

## Author contributions

C.W. and G.B.S. conceived the project. S.R.L. and C.W. designed the study. S.R.L., N.H.C. and C.W. carried out all experiments. S.R.L. and C.W. analyzed the data. S.R.L., C.W., E.J.H, and G.B.S. wrote the manuscript.

## Conflict of Interest

The authors declare no competing interests

## Notes

https://doi.org/10.5281/zenodo.10733846

### Competing Interest Statement

The authors have declared no competing interest.

### Summary of Updates

Since the previous review, we have conducted careful analyses to improve the manuscript, including the addition of sham controls to validate the specificity of observed sensory deficits, refined statistical testing to confirm significant changes in sensory performance, and detailed correlation analyses linking lesion extent to behavioral outcomes. Furthermore, data presentation has been improved to better distinguish sensory deficits from motor-related effects, ensuring the clarity of our findings.

## References

Aeed, Fadi, Nathan Cermak, Jackie Schiller, and Yitzhak Schiller. 2021. “Intrinsic Disruption of the M1 Cortical Network in a Mouse Model of Parkinson’s Disease.” Movement Disorders: Official Journal of the Movement Disorder Society 36(7): 1565–77. doi:10.1002/mds.28538.

Agid, Yves, France Javoy, Jacques Glowinski, Dominique Bouvet, and Constantino Sotelo. 1973. “Injection of 6-Hydroxydopamine into the Substantia Nigra of the Rat. II. Diffusion and Specificity.” Brain Research 58(2): 291–301. doi:10.1016/0006-8993(73)90002-4.

Aiello, Marilena, Roberto Eleopra, and Raffella I. Rumiati. 2015. “Body Weight and Food Intake in Parkinson’s Disease. A Review of the Association to Non-Motor Symptoms.” Appetite 84: 204–11. doi:10.1016/j.appet.2014.10.011.

Antony, Paul M. A., Nico J. Diederich, and Rudi Balling. 2011. “Parkinson’s Disease Mouse Models in Translational Research.” Mammalian Genome 22(7–8): 401–19. doi:10.1007/s00335-011-9330-x.

Arboretti Giancristofaro, R., Stefano Bonnini, and F. Pesarin. 2006. “Permutation Tests for Heterogeneity Comparisons in Presence of Categorical Variables with Application to University Evaluation.” In SVN. https://sfera.unife.it/handle/11392/523739 (November 19, 2024).

Bagga, V., S.B. Dunnett, and R.A. Fricker. 2015. “The 6-OHDA Mouse Model of Parkinson’s Disease – Terminal Striatal Lesions Provide a Superior Measure of Neuronal Loss and Replacement than Median Forebrain Bundle Lesions.” Behavioural Brain Research 288: 107–17. doi:10.1016/j.bbr.2015.03.058.

Barata-Antunes, Sandra, Fábio G. Teixeira, Bárbara Mendes-Pinheiro, Ana V. Domingues, Helena Vilaça-Faria, Ana Marote, Deolinda Silva, Rui A. Sousa, and António J. Salgado. 2020. “Impact of Aging on the 6-OHDA-Induced Rat Model of Parkinson’s Disease.” International Journal of Molecular Sciences 21(10): 3459. doi:10.3390/ijms21103459.

Bonnini, Stefano, L. Salmaso, and A. Solari. 2005. “Multivariate Permutation Tests for Evaluating Effectiveness of Universities through the Analysis of Students Dropouts.” https://sfera.unife.it/handle/11392/523715?mode=full (November 19, 2024).

Branchi, Igor, Ivana D’Andrea, Monica Armida, Tommaso Cassano, Antonella Pèzzola, Rosa Luisa Potenza, Maria Grazia Morgese, Patrizia Popoli, and Enrico Alleva. 2008. “Nonmotor Symptoms in Parkinson’s Disease: Investigating Early-phase Onset of Behavioral Dysfunction in the 6-hydroxydopamine-lesioned Rat Model.” Journal of Neuroscience Research 86(9): 2050–61. doi:10.1002/jnr.21642.

Brim, Bianca, and Walter Struhal. 2021. “Thermoregulatory Dysfunctions in Idiopathic Parkinson’s Disease.” In *International Review of Movement Disorders*, Elsevier, 285–98. doi:10.1016/bs.irmvd.2021.08.009.

Brooks, Simon P., and Stephen B. Dunnett. 2009. “Tests to Assess Motor Phenotype in Mice: A User’s Guide.” Nature Reviews Neuroscience 10(7): 519–29. doi:10.1038/nrn2652.

Chagas, André M., Lucas Theis, Biswa Sengupta, Maik C. Stüttgen, Matthias Bethge, and Cornelius Schwarz. 2013. “Functional Analysis of Ultra High Information Rates Conveyed by Rat Vibrissal Primary Afferents.” Frontiers in Neural Circuits 7. doi:10.3389/fncir.2013.00190.

Cumming, Kirsten, Angus D. Macleod, Phyo K. Myint, and Carl E. Counsell. 2017. “Early Weight Loss in Parkinsonism Predicts Poor Outcomes: Evidence from an Incident Cohort Study.” Neurology 89(22): 2254–61. doi:10.1212/WNL.0000000000004691.

Francardo, Veronica, Alessandra Recchia, Nataljia Popovic, Daniel Andersson, Hans Nissbrandt, and M. Angela Cenci. 2011. “Impact of the Lesion Procedure on the Profiles of Motor Impairment and Molecular Responsiveness to L-DOPA in the 6-Hydroxydopamine Mouse Model of Parkinson’s Disease.” Neurobiology of Disease 42(3): 327–40. doi:10.1016/j.nbd.2011.01.024.

Frankin, Keith B.J., and George Paxinos. 2008. The Mouse Brain in Stereotaxic Coordinates (2008). 3rd ed.

Frund, I., N. V. Haenel, and F. A. Wichmann. 2011. “Inference for Psychometric Functions in the Presence of Nonstationary Behavior.” Journal of Vision 11(6): 16–16. doi:10.1167/11.6.16.

Gage, Gregory J., Daryl R. Kipke, and William Shain. 2012. “Whole Animal Perfusion Fixation for Rodents.” Journal of Visualized Experiments (65): 3564. doi:10.3791/3564.

Giancristofaro, R. A., M. Bolzan, F. Campigotto, L. Corain, and L. Salmaso. 2010. “Combination-Based Permutation Testing in Survival Analysis.” https://www.semanticscholar.org/paper/Combination-based-permutation-testing-in-survival-Giancristofaro-Bolzan/afb8adcf34f52be07f48f05bb45df0be5cdf0c0b (November 19, 2024).

Green, David M., and John A. Swets. 1966. Signal Detection Theory and Psychophysics. Oxford, England: John Wiley.

Iancu, Ruxandra, Paul Mohapel, Patrik Brundin, and Gesine Paul. 2005. “Behavioral Characterization of a Unilateral 6-OHDA-Lesion Model of Parkinson’s Disease in Mice.” Behavioural Brain Research 162(1): 1–10. doi:10.1016/j.bbr.2005.02.023.

Inagaki, Hidehiko K., Miho Inagaki, Sandro Romani, and Karel Svoboda. 2018. “Low-Dimensional and Monotonic Preparatory Activity in Mouse Anterior Lateral Motor Cortex.” The Journal of Neuroscience 38(17): 4163–85. doi:10.1523/JNEUROSCI.3152-17.2018.

Jafari, Zahra, Bryan E. Kolb, and Majid H. Mohajerani. 2020. “Auditory Dysfunction in Parkinson’s Disease.” Movement Disorders 35(4): 537–50. doi:10.1002/mds.28000.

José Luvizutto, Gustavo, Thanielle Souza Silva Brito, Eduardo De Moura Neto, and Luciane Aparecida Pascucci Sande De Souza. 2020. “Altered Visual and Proprioceptive Spatial Perception in Individuals with Parkinson’s Disease.” Perceptual and Motor Skills 127(1): 98–112. doi:10.1177/0031512519880421.

Juri, Carlos, MariCruz Rodriguez-Oroz, and Jose A. Obeso. 2010. “The Pathophysiological Basis of Sensory Disturbances in Parkinson’s Disease.” Journal of the Neurological Sciences 289(1–2): 60–65. doi:10.1016/j.jns.2009.08.018.

Kesayan, Tigran, Damon G. Lamb, Adam D. Falchook, John B. Williamson, Liliana Salazar, Irene A. Malaty, Nikolaus R. McFarland, et al. 2015. “Abnormal Tactile Pressure Perception in Parkinson’s Disease.” Journal of Clinical and Experimental Neuropsychology 37(8): 808–15. doi:10.1080/13803395.2015.1060951.

Ketzef, Maya, Giada Spigolon, Yvonne Johansson, Alessandra Bonito-Oliva, Gilberto Fisone, and Gilad Silberberg. 2017. “Dopamine Depletion Impairs Bilateral Sensory Processing in the Striatum in a Pathway-Dependent Manner.” Neuron 94(4): 855–865.e5. doi:10.1016/j.neuron.2017.05.004.

Konczak, Jürgen, Alessandra Sciutti, Laura Avanzino, Valentina Squeri, Monica Gori, Lorenzo Masia, Giovanni Abbruzzese, and Giulio Sandini. 2012. “Parkinson’s Disease Accelerates Age-Related Decline in Haptic Perception by Altering Somatosensory Integration.” Brain 135(11): 3371–79. doi:10.1093/brain/aws265.

Lee, Christian R., Alex J. Yonk, Joost Wiskerke, Kenneth G. Paradiso, James M. Tepper, and David J. Margolis. 2019. “Opposing Influence of Sensory and Motor Cortical Input on Striatal Circuitry and Choice Behavior.” Current biology : CB 29(8): 1313–1323.e5. doi:10.1016/j.cub.2019.03.028.

Lundblad, M., B. Picconi, H. Lindgren, and M.A. Cenci. 2004. “A Model of L-DOPA-Induced Dyskinesia in 6-Hydroxydopamine Lesioned Mice: Relation to Motor and Cellular Parameters of Nigrostriatal Function.” Neurobiology of Disease 16(1): 110–23. doi:10.1016/j.nbd.2004.01.007.

Ma, Kai, Nian Xiong, Yan Shen, Chao Han, Ling Liu, Guoxin Zhang, Luxi Wang, et al. 2018. “Weight Loss and Malnutrition in Patients with Parkinson’s Disease: Current Knowledge and Future Prospects.” Frontiers in Aging Neuroscience 10: 1. doi:10.3389/fnagi.2018.00001.

Masini, Débora, Carina Plewnia, Maëlle Bertho, Nicolas Scalbert, Vittorio Caggiano, and Gilberto Fisone. 2021. “A Guide to the Generation of a 6-Hydroxydopamine Mouse Model of Parkinson’s Disease for the Study of Non-Motor Symptoms.” Biomedicines 9(6): 598. doi:10.3390/biomedicines9060598.

Moss, Morgane M., Peter Zatka-Haas, Kenneth D. Harris, Matteo Carandini, and Armin Lak. 2021. “Dopamine Axons in Dorsal Striatum Encode Contralateral Visual Stimuli and Choices.” Journal of Neuroscience 41(34): 7197–7205. doi:10.1523/JNEUROSCI.0490-21.2021.

Ollerenshaw, Douglas R., Bilal A. Bari, Daniel C. Millard, Lauren E. Orr, Qi Wang, and Garrett B. Stanley. 2012. “Detection of Tactile Inputs in the Rat Vibrissa Pathway.” Journal of Neurophysiology 108(2): 479–90. doi:10.1152/jn.00004.2012.

Oppo, Valentina, Marta Melis, Melania Melis, Iole Tomassini Barbarossa, and Giovanni Cossu. 2020. “‘Smelling and Tasting’ Parkinson’s Disease: Using Senses to Improve the Knowledge of the Disease.” Frontiers in Aging Neuroscience 12: 43. doi:10.3389/fnagi.2020.00043.

Pesarin, Fortunato, and Luigi Salmaso. 2010. “The Permutation Testing Approach: A Review.” Statistica 70(4): 481–509. doi:10.6092/issn.1973-2201/3599.

Petersen, Carl C. H. 2007. “The Functional Organization of the Barrel Cortex.” Neuron 56(2): 339–55. doi:10.1016/j.neuron.2007.09.017.

Pont-Sunyer, Claustre, Anna Hotter, Carles Gaig, Klaus Seppi, Yaroslau Compta, Regina Katzenschlager, Natalia Mas, et al. 2015. “The Onset of Nonmotor Symptomsin Parkinson’s Disease (the ONSET PD Study).” Movement Disorders 30(2): 229–37. doi:10.1002/mds.26077.

Prätorius, B, S Kimmeskamp, and T.L Milani. 2003. “The Sensitivity of the Sole of the Foot in Patients with Morbus Parkinson.” Neuroscience Letters 346(3): 173–76. doi:10.1016/S0304-3940(03)00582-2.

Przedborski, S, and V Jackson-Lewis. 1995. “DOSE-DEPENDENT LESIONS OF THE DOPAMINERGIC NIGROSTRIATAL PATHWAY INDUCED BY INTRASTRIATAL INJECTION OF 6-HYDROXYDOPAMINE.” Neuroscience 67(3): 631–47.

Quiroga-Varela, A., E. Aguilar, E. Iglesias, J.A. Obeso, and C. Marin. 2017. “Short- and Long-Term Effects Induced by Repeated 6-OHDA Intraventricular Administration: A New Progressive and Bilateral Rodent Model of Parkinson’s Disease.” Neuroscience 361: 144–56. doi:10.1016/j.neuroscience.2017.08.017.

Richardson, Kelly C., and Joan E. Sussman. 2019. “Intensity Resolution in Individuals With Parkinson’s Disease: Sensory and Auditory Memory Limitations.” *Journal of Speech*, Language, and Hearing Research 62(9): 3564–81. doi:10.1044/2019_JSLHR-H-18-0424.

Ritt, Jason T., Mark L. Andermann, and Christopher I. Moore. 2008. “Embodied Information Processing: Vibrissa Mechanics and Texture Features Shape Micromotions in Actively Sensing Rats.” Neuron 57(4): 599–613. doi:10.1016/j.neuron.2007.12.024.

Stüttgen, Maik C., Johannes Rüter, and Cornelius Schwarz. 2006. “Two Psychophysical Channels of Whisker Deflection in Rats Align with Two Neuronal Classes of Primary Afferents.” The Journal of Neuroscience 26(30): 7933–41. doi:10.1523/JNEUROSCI.1864-06.2006.

Thiele, Sherri L., Ruth Warre, and Joanne E. Nash. 2012. “Development of a Unilaterally-Lesioned 6-OHDA Mouse Model of Parkinson’s Disease.” Journal of Visualized Experiments (60): 3234. doi:10.3791/3234.

Tieu, K. 2011. “A Guide to Neurotoxic Animal Models of Parkinson’s Disease.” Cold Spring Harbor Perspectives in Medicine 1(1): a009316–a009316. doi:10.1101/cshperspect.a009316.

de la Torre-Martinez, Roberto, Maya Ketzef, and Gilad Silberberg. 2023. “Ongoing Movement Controls Sensory Integration in the Dorsolateral Striatum.” Nature Communications 14(1): 1004. doi:10.1038/s41467-023-36648-0.

Ungerstedt, Urban, and Gordon W. Arbuthnott. 1970. “Quantitative Recording of Rotational Behavior in Rats after 6-Hydroxy-Dopamine Lesions of the Nigrostriatal Dopamine System.” Brain Research 24(3): 485–93. doi:10.1016/0006-8993(70)90187-3.

Waiblinger, Christian et al. 2022. “Emerging Experience-Dependent Dynamics in Primary Somatosensory Cortex Reflect Behavioral Adaptation.” Nature Communications 13(1): 534.

Waiblinger, Christian, Clarissa J. Whitmire, Audrey Sederberg, Garrett B. Stanley, and Cornelius Schwarz. 2018. “Primary Tactile Thalamus Spiking Reflects Cognitive Signals.” The Journal of Neuroscience 38(21): 4870–85. doi:10.1523/JNEUROSCI.2403-17.2018.

Waiblinger, Christian et al. 2019. “Stimulus Context and Reward Contingency Induce Behavioral Adaptation in a Rodent Tactile Detection Task.” The Journal of Neuroscience 39(6): 1088–99.

Wichmann, Felix A., and N. Jeremy Hill. 2001. “The Psychometric Function: I. Fitting, Sampling, and Goodness of Fit.” Perception & Psychophysics 63(8): 1293–1313. doi:10.3758/BF03194544.

Willis, A. W., E. Roberts, J. C. Beck, B. Fiske, W. Ross, R. Savica, S. K. Van Den Eeden, C. M. Tanner, and C. Marras. 2022. “Incidence of Parkinson Disease in North America.” npj Parkinson’s Disease 8(1): 1–7. doi:10.1038/s41531-022-00410-y.

Wolfe, Jason, Dan N Hill, Sohrab Pahlavan, Patrick J Drew, David Kleinfeld, and Daniel E Feldman. 2008. “Texture Coding in the Rat Whisker System: Slip-Stick Versus Differential Resonance” ed. Garrett B Stanley. PLoS Biology 6(8): e215. doi:10.1371/journal.pbio.0060215.

