## Supplementary material for "The 6-OHDA Parkinson’s Disease Mouse Model Shows Deficits in Sensory Behavior": Figure 3-1. Extended data supporting Figure 3.

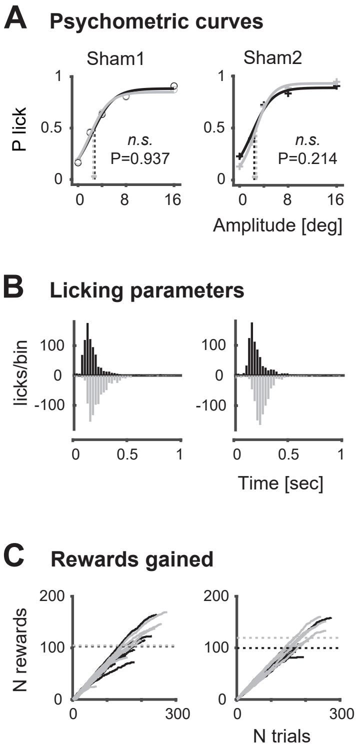


**Figure 3-1. Extended data supporting Figure 3. Assessment of sensory capabilities in sham controls. A.** Psychometric curves and response thresholds for two control animals performing the detection task before (black) and after sham injection (gray). Each data point represents an animal's response probabilities at a given stimulus amplitude, averaged across sessions. Solid curves are logistic fits to the data. Response thresholds at P = 0.5 are shown as vertical lines with 95% confidence limits at the bottom. A non-parametric permutation test combined with Fisher's method was used to compare each subject's psychometric data before and after sham lesion (*n.s.*, P > 0.05). **B.** Histograms display licks/bin (25 ms bins) for individual mice before and after sham lesion. **C.** Number of rewards accumulated across trials by each animal before and after sham lesion. Each line represents a single session, with dashed horizontal lines indicating the average total rewards per session.
