## Supplementary material for "The 6-OHDA Parkinson’s Disease Mouse Model Shows Deficits in Sensory Behavior": Figure 4-1. Extended data supporting Figure 4.

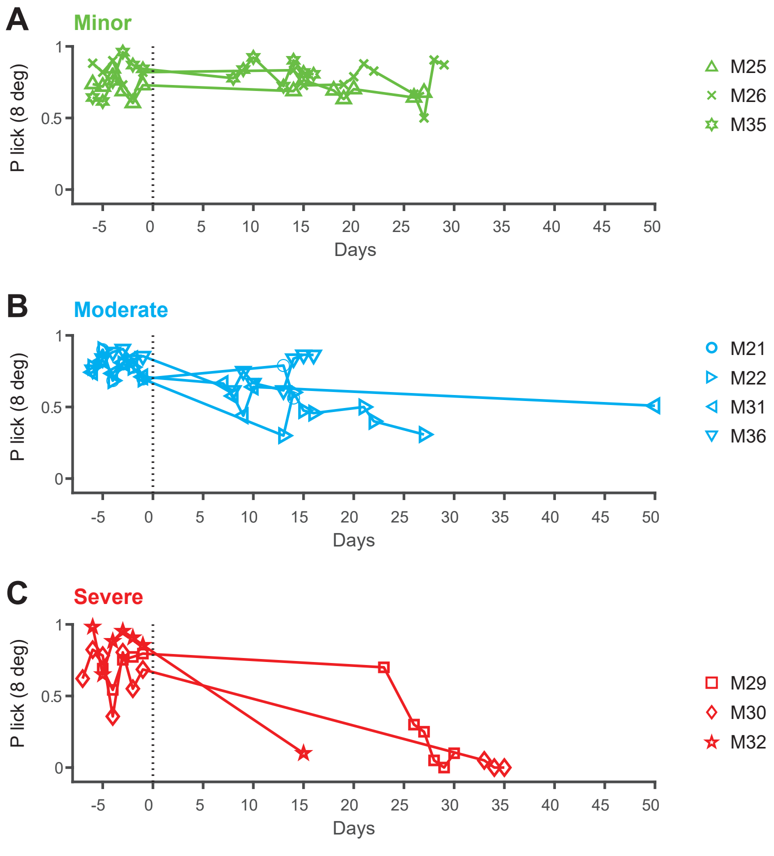


**Figure 4-1. Extended data supporting Figure 4. Detection performance over days.** The 3 categories, minor, moderate and severe, are based on the difference in thresholds from the psychometric curves (see Figure 3). **A.** Detection performance (P correct lick with 8 degree whisker deflection) over time (days) for n=3 animals with minor deficits, before and after the 6-OHDA lesion. **B.** Performance for n=4 animals with moderate deficits. **C.** Performance for n=3 animals with severe deficits. Testing was paused during recovery periods (no data points).
